# Error correction and assembly complexity of single molecule sequencing reads

**DOI:** 10.1101/006395

**Authors:** Hayan Lee, James Gurtowski, Shinjae Yoo, Shoshana Marcus, W. Richard McCombie, Michael Schatz

## Abstract

Third generation single molecule sequencing technology is poised to revolutionize genomics by enabling the sequencing of long, individual molecules of DNA and RNA. These technologies now routinely produce reads exceeding 5,000 basepairs, and can achieve reads as long as 50,000 basepairs. Here we evaluate the limits of single molecule sequencing by assessing the impact of long read sequencing in the assembly of the human genome and 25 other important genomes across the tree of life. From this, we develop a new data-driven model using support vector regression that can accurately predict assembly performance. We also present a novel hybrid error correction algorithm for long PacBio sequencing reads that uses pre-assembled Illumina sequences for the error correction. We apply it several prokaryotic and eukaryotic genomes, and show it can achieve near-perfect assemblies of small genomes (*<* 100Mbp) and substantially improved assemblies of larger ones. All source code and the assembly model are available open-source.

## 1 Introduction

Three new 3^rd^ generation single molecule sequencing technologies are currently available from Pacific Biosciences (PacBio) [Roberts *et al.*, 2013], Moleculo [Voskoboynik *et al.*, 2013], and Oxford Nanopore [Hayden, 2014]. The most established of these is the Single Molecule Real Time (SMRT) sequencing platform produced by PacBio. Their current instrument, the PacBio RS II, can generate reads as long as 54kb with an average read length over 10kbp, approximately 50 to 250 times longer than those available from the widely used 2^nd^ generation Illumina platform [56] (Figure 1). The technology uses a powerful imaging system, called a zero-mode waveguide, to image fluorescently tagged nucleotides as they are incorporated along individual template molecules. Alternatively, Moleculo uses dilution, clonal amplification, and barcoding to sequence long molecules of DNA of up to 8 to 10kbp [Voskoboynik *et al.*, 2013], and Oxford Nanopore detects changes to ion flow as nucleotides pass through a nanopore, achieving reads as long as 10kbp in prototype instruments [Hayden, 2014].

**Figure 1:**
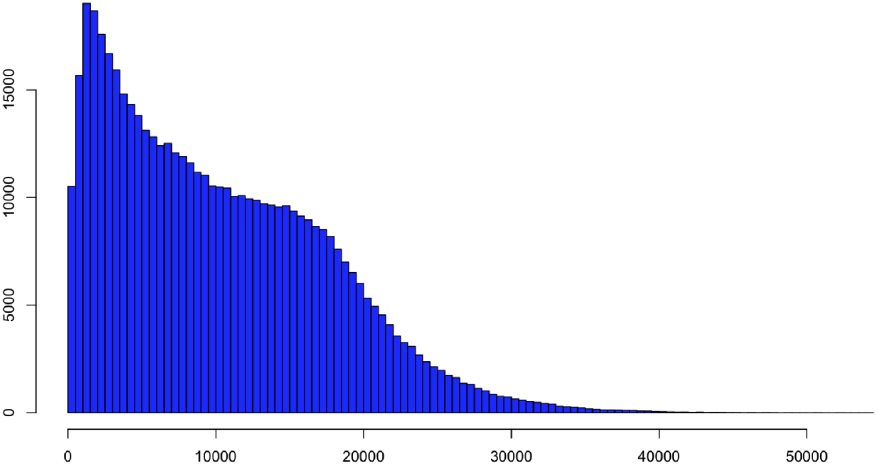
PacBio read length distribution for the recently sequenced rice genome (IR64) using C3-P5 chemistry sequenced at PacBio. The mean read length was 10,232bp, and the maximum extends to 54,288bp.

The long reads produced by these instruments are an enabling technology for a wide variety of important genomics applications: in *de novo* genome assembly, the longer reads span more repetitive elements making it possible to assemble more contiguous sequences, up to the assembling complete chromosomes directly from the shotgun sequences. (Figure 2). In genome resequencing, Moleculo reads have been used to phase haplotypes [Kuleshov *et al.*, 2014], and long PacBio reads have been successfully used to resolve complex structural variations [Maron *et al.*, 2013]. In transcriptome analysis, the long reads span more exon junctions, or even entire transcripts, making it possible to resolve individual isoforms of complex genes [Sharon *et al.*, 2013].

**Figure 2:**
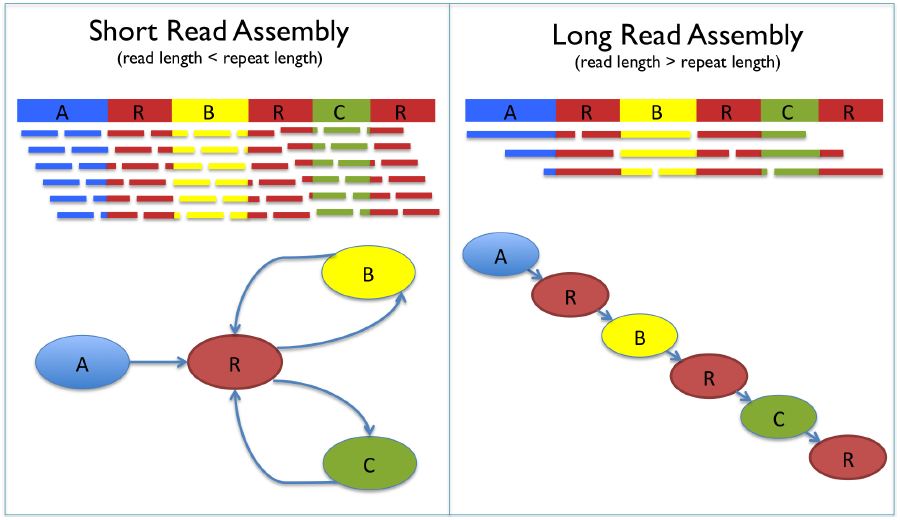
Advantages of long read assembly. Shown here is a genome sequence with 3 unique segments A, B, and C in blue, yellow, and green, and 3 copies of a repeat R in red. If the read length is shorter than the repeat region (left), the corresponding assembly graph will contain 4 contigs, with edges connecting them but in an ambiguous order. If the reads span the repeat segments (right), then the corresponding assembly graph will not be branching, and a single long contig spanning the entire genome will be created.

In a *de novo* genome assembly, the reads are compared to each other to construct an assembly graph, such as the widely used string graph [Myers, 2005] or de Bruijn graph [Kingsford *et al.*, 2010]. The graph is then simplified and traversed to reconstruct larger sequences, ideally assembling entire chromosomes into individual contigs. The assembly graph, and the overall quality of the assembly, is limited by several technical and biological reasons. At low coverage, contigs can end due to gaps of coverage or errors, but more significantly, even at high coverage repeats longer than the read length lead to “branches” in the assembly graph, effectively ending the contigs. Longer read lengths are thus highly advantageous as they will be able to span across more repeats than short reads. The potential for more uniform coverage from single molecule sequencing also minimize the coverage gaps that can occur with short read sequencing [Ross *et al.*, 2013].

The most widely used statistical model of genome assembly was developed by E.S. Lander and M.S. Waterman in 1988 ([Lander and Waterman, 1988]) based on a Poisson coverage distribution. For the last 25 years, this simple model has guided researchers and lead to useful recommendations about minimum coverage and other guidelines. However it serves as only a crude guide, and can also predict nonsensical results such as with 100x coverage of 100bp reads, the human genome should assemble into contigs hundreds of gigabases long, far beyond the length of the genome itself (Supplementary Figure S1). In contrast, the best de novo genome assemblies with these data have had average contig sizes of only 20kbp to 30kbp [Gnerre *et al.*, 2011; Simpson *et al.*, 2009]).

The prodigious lack of predictive power is because the Lander-Waterman model assumes the genome is free of any significant repetitive sequences. In real genomes, however, repeats are ubiquitous, and genome assemblers end contigs at their boundaries if not spanned by sufficiently long reads. Interestingly, a modest increase of read length can exert a significant improvement to the assembly performance. This is because the repeat distribution in a typical eukaryotic genome is exponentially decreasing (Supplementary Figure S3). Consequently, in the rice genome, increasing the read length 30 fold from a typical Illumina read length (100bp) to a typical PacBio read length (3650bp) exponentially decreases the number of repeats that are not fully spanned by more than 300 fold. Previous work modeling repeats in an assembly [Kingsford *et al.*, 2010] has focused on exact repeats, but this is an unrealistic assumption and requires perfect sequence fidelity. Instead, in practice assemblers evaluate the rate of differences, and accept a low rate of differences (2%-3%) as sequencing errors [Koren *et al.*, 2012; Miller *et al.*, 2008]. As a result, an assembler can only reliably resolve identical or near identical repeats if there are reads that completely span the repeat and into the flanking unique sequences.

To build a realistic model of assembly that accounts for the complexities of real genomes, we adopted a datadriven approach using Support Vector Regression (SVR). For this, we mimic current sequencing technology as much as possible and simulate reads from 26 reference genomes at 4 coverage levels and 5 read lengths using realistic read length distributions from PacBio instruments. Then we use an enhanced version of the Celera Assembler [Koren *et al.*, 2012; Miller *et al.*, 2008] to assemble the simulated reads. In this way the assembler informs us of what can be assembled and which repeats are too complex. The resultant model accurately predicts genome assembly as a function of read length and coverage, and in leave-one-species-out cross validation is within 10% accurate of residual boundary.

In building this data driven model, it was natural to compare some existing de novo assemblies to the theoretical upper bounds generated by our model. Assemblies produced with PacBio’s SMRT sequencing data were in some cases less contiguous than predicted. Ultimately the pre-assembly error correction of the PacBio data was at fault, and we began looking at ways to better assembly contiguity through an improved error correction pipeline.

The main hurdle in using PacBio’s long reads (PBLR) in de novo assembly is the high per-base error rate of approximately 15% [Chin *et al.*, 2013]. There are no assemblers that are currently equipped for this high of an error rate natively. Instead read correction pipelines have been developed to address this problem. Koren et al developed a method called PacBioToCA using the Celera Assembler’s overlap machinery to correct PBLRs using high-identity short read data produced from the same sample [Koren *et al.*, 2012]. A conceptually similar approach, HGAP [Chin *et al.*, 2013] - developed by Pacific Biosciences, does not require a second high-identity library, but instead relies on high PBLR coverage to overcome the high error rate. HGAP and PacBioToCA perform very well on genomes where high PBLR coverage can be obtained: the accuracy of the error corrected reads approaches or exceeds 99%, and the accuracy of the final consensus can exceed 99.999% [Chin *et al.*, 2013]. However, when used for larger genomes, these tools begin to falter, in particular, because HGAP and the new self-correction mode of PacBioToCA require very high PBLR coverage (*>* 50x), and it can be prohibitively expensive to obtain the necessary coverage.

In response to these challenges, we developed an improved error correction pipeline that uses pre-assembled high-accuracy contigs from inexpensive short read data as the basis of the correction. In our testing of several prokaryotic and eukaryotic genomes we find that this approach can greatly outperform both the older PacBioToCA pipeline and HGAP pipeline at intermediate long read coverage levels (5-50x coverage). Using this approach, we have built perfect assemblies of *E. coli* and near perfect assemblies of yeast, in addition to greatly improved assemblies of *Arabidopsis thaliana* and rice; all approaching the optimal value predicted by the model.

All of the source code, sequencing reads, assemblies, and assembler parameters used in this study are available online at http://schatzlab.cshl.edu/data/ectools/. The open source correction pipeline and manual are available at https://github.com/jgurtowski/ectools/. The prediction model available online as an interactive webapp at http://qb.cshl.edu/asmmodel/predict.html (Supplementary Figure S9) as a resource to guide all future genome assembly projects.

## 2 RESULTS

### 2.1 Assembly Performance

Several important considerations must be taken into account when sequencing a genome: How much coverage do we need? How long should we expect the contigs to be given a certain read length and coverage?; How long should the reads be to assemble into one contig per chromosome?; To answer these questions, we systematically analyzed 26 genomes ranging in size from the 1.66 Mbp *M. jannaschii* genome to the 3.0 Gbp *H. sapiens* genome (Table S1). These genomes were selected to be a diverse, representative sample of genomes across the tree of life, consisting of 5 bacteria, 1 archae, 3 fungi, 1 amoebazoa, 8 plants, 3 invertebrates and 5 vertebrates species. Whenever multiple genomes of similar size were available, we selected the genome with the highest quality sequence to ensure the analysis best captures the true complexities present. Notably, we excluded the largest currently available genomes, such as the 22 Gbp Norway spruce [Nystedt *et al.*, 2013], since the N50 sizes of these assemblies are unrealistically low (*<* 50kbp), and would have distorted the analysis of the repeats present.

After selecting the reference genomes, we simulated shotgun sequencing them with a variety of read lengths and coverage values. We began by simulating reads with the “C2” PacBio read length distribution, with a mean read length of 3,650bp (mean1) and their newer “C3” chemistry which has a mean read length approximately double this (7,400bp, mean2). Since PacBio sequencing has historically doubled their average read length every six months to one year, we also considered the effects of doubling the mean a total of five times (120Kbp, mean32) to project the technologies that should become available over the next few years. For each of those read length distributions, we simulated 5x to 40x coverage of the genomes, and assembled the reads using an enhanced version of the Celera Assembler that supports reads up to 512kbp. Given the rapid improvement to sequencing technologies and especially the error correction algorithms that can post-process the reads into virtually perfect reads, we used error free reads for the simulations to establish an upper-bound on the assembly quality and focus on the underlying biological complexities. However, the assembler will still compare the reads using standard parameters, so that inexact repeats within 3% similarity that are longer than the read lengths will remain unresolved.

To quantify and normalize assembly quality for different genome sizes and numbers of chromosomes, we define assembly performance as follows:

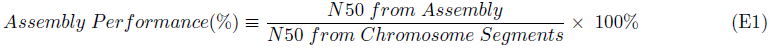

A chromosome segment is a sequence of a chromosome free of extended runs of 1 million or more consecutive “N”s (other Ns in the genome are converted into A’s to transform it into a repeat of maximal complexity). For most species, chromosome segments are equivalent to chromosomes and assembly performance measures the achievable contig N50 compared to the idealized N50 size of the chromosomes. However, some species, including human, mouse, and lizard, have especially long runs of Ns in their centromeric or telomeric regions that would have substantially decreased the possible assembly performance. Evaluating the chromosome segments gives a more realistic estimate of assembly performance of the euchromatic regions, and effectively evaluates the performance relative to the sizes of the chromosome arms.

The results of the assemblies, and a selection of genuine long read assembly results, are shown in Figure 3. Assembly performance follows an overall logistic shape: the performance is consistently high for small genomes, and drops off as the genome size increases. The rate the performance drops off depends on the read length distribution, but longer reads consistently improve assembly performance until the genome is fully assembled (Supplementary Figure S2). It is notable that with the current PacBio read length distributions (C2 & C3 / mean1 & mean2), the assembly performance for most genomes less than 100Mbp is near 100%, meaning it should be possible to assemble the complete chromosomes of many of these species using the currently available technology. Indeed in our own testing we have achieved perfect or near-perfect assemblies of several microbial genomes and lower eukaryotic genomes. Beyond this size, the currently available read lengths substantially improve the assembly compared to typical short read assemblies, although the achievable performance is still below entire chromosome segments. For example, using current mean1 or mean2 reads, the assembly performance of the human genome is only 12% or 28% while reads 32 times longer (mean32) are needed to assemble it completely.

**Figure 3:**
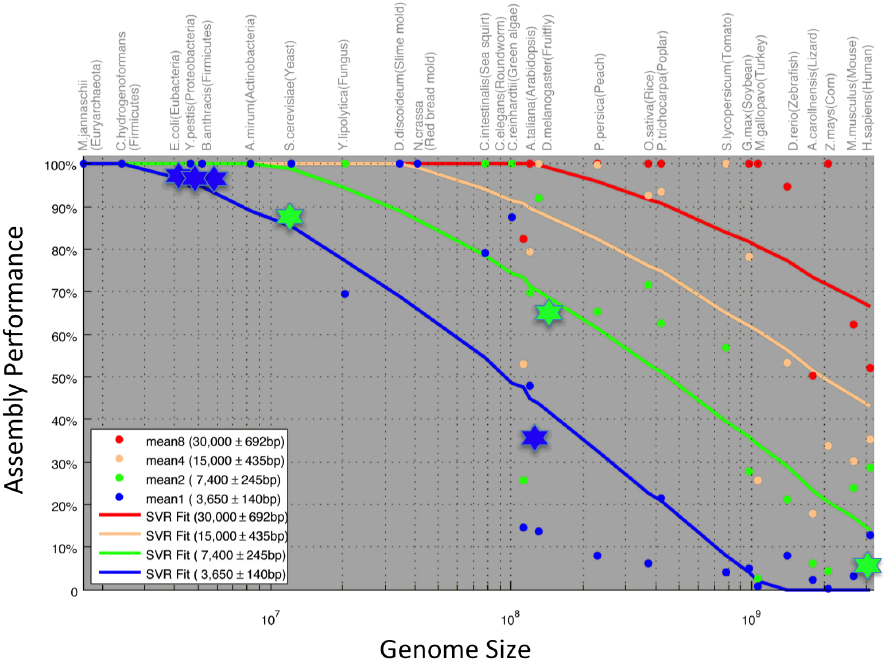
Assembly performance. The x-axis measures the genome size of the 26 genomes in log space. The y-axis measures the assembly performance of the different assemblies using mean1 through mean4 reads. Points indicate the results of simulated experiments with 20x coverage. Stars indicate the results of the assembly of real genomes. Lines show the best fit line from the SVR model. See Supplementary Table 1 for the values of the simulated datasets and supplementary Table S5 for the genuine data sets.

### 2.2 Genome Assembly Modeling

From these data and assuming the read accuracy will be sufficiently high through improved sequencing or error correction technologies, we determined assembly performance primarily depends on four major features: mean read length (L), coverage (C), genome size (G), and repeat complexity (R) as we model in (2). Here, we define repeat complexity (R) as the number of repeats longer than the mean read length L.

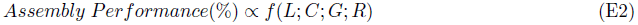

For a given read length distribution, higher coverage is consistently correlated with improved assembly performance although the gains diminish beyond 20x coverage (Supplementary Figure S2). At very low coverage, contigs often end because of coverage gaps. By 20x coverage, though, with high probability every base in the genome should be sequenced, so contig breaks will primarily be due to repeats that are not fully spanned by long reads. For example, when the coverage of the human genome increases from 5x to 20x, the assembly performance improves significantly, from a contig N50 of just 41.6Kbp to 11.5Mbp for mean1 and from 5.2Mbp to 80.9Mbp for mean32. Beyond 20x coverage, the additional gains are more modest, although some improvement is possible by progressively spanning more of the repeats, especially those whose lengths are near the maximum read length in the distribution. In the human assemblies, increasing the coverage from 20x to 40x improves the contig N50 from 11.5Mbp to 22.5Mbp for the mean1 assembly and from 80.9Mbp to 86.7Mbp for the mean32 assembly.

Repeat complexity is more difficult to model since it is external to the characteristics of the sequencing technology, and in principle two genomes of similar size could have different complexities and assembly performances. Indeed, *A. thaliana*(120Mbp) and *D. melanogaster* (130Mbp), have similar size but *D. melanogaster* has many more repeats longer than 3,600bp (mean1) than *A. thaliana* leading to a superior assembly for *A. thaliana*. At 30,000bp (mean8), however, the complexities reverse, and the assembly performance of *A. thaliana* is worse than *D. melanogaster*. However, over broad genomic size scales there is nevertheless a strong linear correlation between genome size and repeat complexity (Supplementary Figure S6). There are also a few outlier species with unexpectedly short maximum repeats and a few with unexpectedly long, although we speculate that the outliers, especially genomes with surprisingly few long repeats, are partially explained by limitations in the technology used to construct them rather than a true biological result.

Using these four features (L, C, G, R), we used support vector regression (SVR) to construct a model of assembly performance (See Online Methods). The results of the final model have strong predictive power: in leave-one-species-out cross validation, the model can predict the performance of given species is within 9% of residual boundary. A simplified model (*γ*=1, C = 100, *ϵ* = 10) is also available as a web service (Supplementary Figure S9) [Lee, 2013], which predicts the assembly performance for a user specified genome length without an explicit value for the repeat complexity. Interestingly, this simplified model has similar accuracy to the full model, so that the webapp can display the read length and coverage tradeoffs for genomes of any user specified size.

### 2.3 Hybrid Error Correction and Assembly

The aforementioned model not only sheds light on the future of assembly performance, it also sets the bar for current assembly projects. Some current assembly projects using real data fell short of the ideal performance predicted by our model. Our investigation revealed that error-rate plays a large part, and in particular PacBioToCA has a non-random bias against PacBio reads with high error rate (Figure 4) and HGAP fails to adequately correct lower coverage datasets. As a result, reads critical to the assembly were being split into short segments or completely lost thus reducing their ability to span repeats and form large contigs.

**Figure 4:**
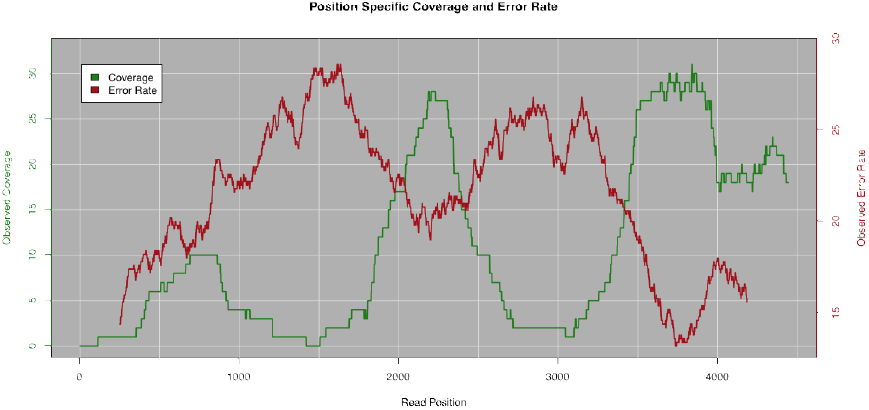
Error rate and PacBioToCA coverage of an individual read. The plot shows the characteristics of an individual PacBio read in the pipeline. The red curve shows the local error rate relative to the reference genome computed by a 200bp sliding window and shows the error rate can fluctuate from 15% to nearly 30%. The green curve shows the number of short reads that could be aligned by the PacBioToCA pipeline at each position in the read. The error rate and coverage levels are anti-correlated, which resulted in the read being split into multiple segments after correction.

To remedy this issue, we developed a new correction pipeline, ECTools, that takes as input a short-read assembly as the backbone for correction (See Online Methods). This approach has the advantage of aligning PacBio reads to pre-assembled contigs that provide more context for seeding alignments as compared to short reads alone. We tested this approach in several assembly projects including *E. coli*, *S. cerevisiae* (yeast), *Arabidposis thaliana* Ler-0, and *Oryza sativa* pv Indica IR64 (rice) for which long read coverage and reference genomes were available. The assembly results can be found in Figure 5 and in supplementary note 1.

**Figure 5:**
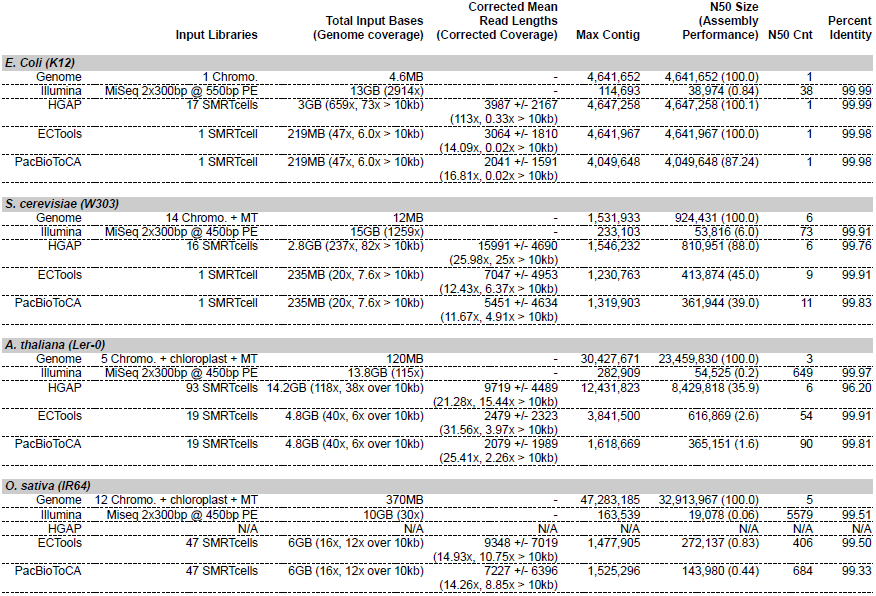
Assembly comparison of four species using three different correction approaches. HGAP is the self-correction technique designed by Pacific Biosciences. PacbioToCA is a hybrid correction approach integrated into the Celera assembler. ECTools is a new hybrid correction algorithm specifically designed for large genomes. See supplementary note for a description of assembly parameters.

Both the *E. coli* and yeast assemblies were used to compare deep-coverage self-correction with HGAP against low-coverage hybrid corrections using both ECTools and PacBioToCA. The deep coverage self correction produces assemblies that are perfect in the case of *E. coli* and nearly perfect, i.e. a single or very small number of contigs per chromosome in yeast. The hybrid assemblies, using just a single SMRTcell, also produce perfect assemblies in *E. coli* and excellent assemblies with only a few additional breaks in yeast. For example, in *S. cerevisiae*, the N50 count (the number of contigs needed to span half the genome) was 6 for self-correction with all of the data and the best ECTools hybrid assembly had an N50 count of 9. These results highlights both the effectiveness of hybrid error correction as well as the decreasing marginal gains from very deep sequencing confirming the results of the simulations.

The same experiment was performed in *A. thaliana* for which substantial C2 long read coverage is available. Because this genome is more complex than both yeast and *E. coli*, there is a larger gap between hybrid and non-hybrid correction performance. However, the amount of sequencing necessary to complete the selfcorrection is quite expensive by today’s standards, and for a genome this large or larger, most projects would only have available what we used in the hybrid assemblies; making them a more useful measure of today’s assembly performance. That said, the assemblies produced by ECTools are quite impressive with N50 of nearly 616kb using only a third of the coverage of the self correction techniques. Furthermore, ECTools begins to substantially outperform PacBioToCA, making it a better choice as genome sizes get larger.

Finally, the rice strain IR64 was sequenced to roughly 16x coverage using the Pacbio RS II with C3 chemistry (Figure 1). This coverage level is too low to run HGAP and no self-correction assembly was generated. However after hybrid error correction with ECTools, the results show that we were able to produce an N50 contig size of 272kb. This is more than one order of magnitude better than the Illumina-only ALLPATHS assembly and more than 5 times better than the contig N50 of the existing reference genome giving new insights into genes, regulatory regions, and structural information not available otherwise.

## 3 Discussion

Researchers are currently reporting that PacBio-based assemblies of microbes can be far better than short-read assemblies, including perfectly assembled chromosomes directly from the shotgun reads [Koren *et al.*, 2013]. As was experienced with 1st and 2nd generation sequencing, it is only a matter of time before results seen in microbes are translated to larger genomes. The most significant limiting factors are the comparatively lower throughput and higher error rate. As such, the most practical way to sequence a large genome is a hybrid approach combining deep coverage of inexpensive short reads with moderate coverage of long reads. Using our improved algorithm, this can lead to excellent assemblies: in the complex rice genome, over 10 times improved contiguity over a state-of-the-art Illumina-only assembly, and nearly 3 times better than older hybrid assembly approaches. The error corrected reads are virtually perfect, and could be used for non-assembly projects, such as mapping structural variations or identifying alternative splicing.

We have also developed a predictive model of assembly performance, using a data-driven approach by assembling simulating reads from published reference genomes. As such it should be considered an upper-bound: the genomes we analyzed have gaps and errors that will mask their true complexity, and our simulated reads did not contain errors nor heterozygosity. Nevertheless the results are useful for analyzing the libraries needed to achieve results as good as those previously published assemblies. In practice, researchers may need to oversample the genome more than predicted to account for any residual errors or biases present.

The model and our own experiences with real data show it should be possible to achieve nearly complete chromosomes for genomes up to 100Mbp in size with the currently available single molecule sequencing. For larger genomes, great gains are possible over short read sequencing, with results approaching or exceeding those from BAC-by-BAC assemblies. If the project demands even higher quality assemblies or complete genomes, the model also forecasts when those data may be available. In particular, for the human genome the read lengths need to double approximately 4 more times before complete chromosomes should be possible. While this seems far off in the future, if the historical trends continue this could be achieved in as little as 3 to 4 years. When that milestone is reached, it is likely that many projects will begin from the fully assembled genomes instead of variant lists. Until then, we are also very excited by the potential for combining these advances with emerging methods for long-range scaffolding, such as by using chromatin conformation ([Burton *et al.*, 2013; Kaplan and Dekker, 2013]). Whereas scaffolding methods are currently limited by the short contigs generated from short read assemblies, when combined with long read long contigs, the methods should be able to reliably scaffold the contigs into complete chromosome arms.

## 4 ONLINE METHODS

### 4.1 Reads Simulation for Model Species

For model fitting and prediction, we simulated PacBio-like reads for 26 species ranging in size from *M.jannaschii* (1.66Mbp) to human (3Gbp). Our simulator, ReadSim, generates long reads mimicking the read length distribution present in an input file by selecting a random starting position in the genome, and generating a read of the next observed length. For the simulations, we used the 1.28M read lengths derived from PacBio “C2” sequencing of the rice Nipponbare genome as our baseline mean1 read length distribution (Figure 6). For other read lengths distributions, we scaled each observed read length by 2 to 32 times, thus shifting the mean read length and standard deviation while maintaining the same overall shape to the distributions. For each species, we simulate four different read lengths (mean1, mean2, mean4, mean8), at four different coverages: 5x, 10x, 20x and 40x. For human, we considered two more mean read length distributions (mean16 and mean32) because shorter reads were not sufficient to completely assemble the chromosomes into single contigs. Overall, 424 samples were assembled: 4 read lengths and 4 coverage levels were used for 25 species, and 6 read lengths at 4 coverage levels were used for the human genome.

**Figure 6:**
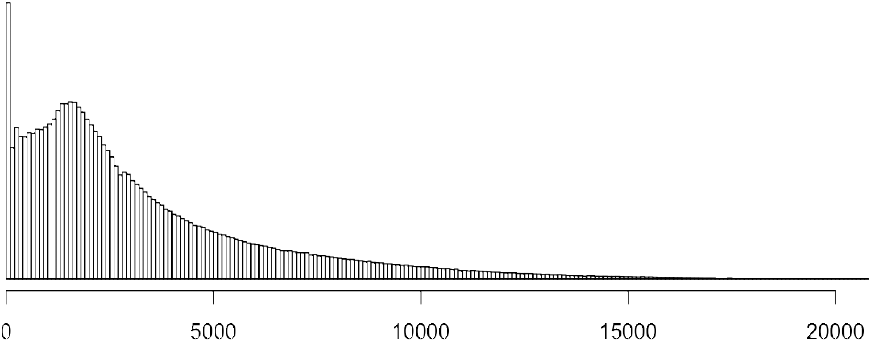
Mean1 Read Length Distribution. Histogram of the read length distribution of rice (Nipponbare) sequenced using PacBio C2-XL chemistry. The mean length is 3290 and maximum length 24,405.

### 4.2 Celera Assembler Enhancement for Long Reads

We previously enhanced the Celera Assembler to assemble reads as long as 32kbp long, but additional modifications were necessary to support the longest reads in this study with a reasonable amount of RAM. First, the compile time option *AS READ MAX NORMAL LEN BITS* was increased from 15 to 19, so that the assembler would store reads as long as 524,288bp. The worker thread stack size in the overlapper was increased from 4MB to 128MB so that each thread had enough space to process long reads in parallel mode. Unnecessary memory allocations were removed and primitive types for some variables, e.g. *MAX ERRORS*, were updated to 64 bits. A new structure, *nm_t_* was added to use 32 bits instead of 16 bits during overlapping. With these modifications all 424 assemblies were computed on the BlackNBlue cluster at Cold Spring Harbor Laboratory, which contains a total of 1,696 cores over 102 nodes. Two high memory nodes with 1.5TB of RAM and 64 cores were used for the most memory intensive stages, especially the unitigger, while the other 100 standard compute nodes with 128 GB of RAM and 16 cores were used to parallelize overlap and the overlap-based error correction stages using Univa Grid Engine. All of the modifications are available open source on the Celera Assembler Website http://wgs-assembler.sf.net.

### 4.3 Feature Engineering

Although assembly performance generally depends on genome size, read length, coverage and repeats, we performed feature engineering to boost predictive power. For example, the correlation coefficient of genome size and read length with assembly performance are 0.38 and 0.2, respectively, but the correlation of *log*(*genomesize*) and *log*(*readlength*) are 0.49 and 0.32, respectively. More significantly, we determined the correlation of *log*(*genome*)*/log*(*readlength*) is 0.6, which we use as the first independent variable in the model. The next independent variables used are *log*(*coverage*), which has an *r*^2^ = 0.58 and *log*(*repeatcount*), which has an *r*^2^ = 0.44.

### 4.4 Support Vector Regression (SVR)

With these three carefully selected variables, we used Support Vector Regression (SVR) to derive a model of their relationships. SVR is one of the widely used machine learning algorithms for modeling and prediction because of (1) its robustness to overfitting/outliers and (2) its generalizability that provides the simplest model given fixed amount of training errors. See Smola and Schölkopf [2004] for a tutorial. Its robustness to overfitting is essential when the sample size is small, and is achieved by *ε*-insensitive linear loss function (E3). With this function, if the difference between true value and predicted value is less than *∈*, the loss is considered zero. If, however, the difference is more than *∈*, the loss is computed as sum of absolute values of their differences. The *∈*-insensitive property forms an area, called the safety boundary, that allows for smoother values and reduces overfitting.

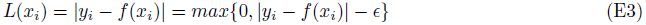

Robustness to outliers is a very important property when sample qualities are variable. This is essential to our analysis, since the reference genomes have errors and omissions in them. Model robustness also dampens the variability in assembly performance that occurs from the stochastic placement of reads where relatively small differences in the assembly may occur at a given coverage level as the exact positions of the reads shift from run to run. The robustness to outliers of SVR is achieved by linearity of loss function. Classic regression uses quadratic form of loss function (E4), in which outliers are weighed heavily by their distance from the learned function. In contrast, the linear loss function of SVR (E3) weighs all points evenly, so that outliers will not influence the parameters as much.

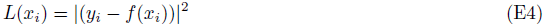

SVR is also well-known for its generalizability. When the optimization process minimizes the *epsilon*-insensitive linear loss function, it naturally pursues a safety area as large as possible. This makes the loss easier to minimize, and consequently, the fitted function will be simple as possible given a fixed amount of training errors.

### 4.5 Model Learning and Grid Search for Parameter Space

We used LIBSVM to compute the SVR, which won many international prediction competitions [Chang and Lin, 2011] for its excellent prediction power and speed. Using LIBSVM, we tested polynomial kernels up to degree of four and Radial Basis Function(RBF) to compute overall performance (Supplementary Figure S7). For the each degree of the polynomial kernels, we performed grid search from 10*^−^*^6^ to 10^6^ for C(option -c), which is a tradeoff between function complexity 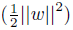 and loss 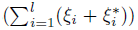 and also plays as a regularization (E5), and from 10*^−^*^6^ to 10^5^ for *∈*(option -p), which decides the safety region around the fit. For the RBF kernel, we searched from 10*^−^*^6^ to 10^3^ for *γ*(option -g), from 10*^−^*^6^ to 10^6^ for C and from 10*^−^*^6^ to 10^5^ for 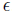.

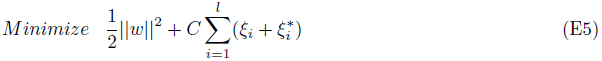

Subject to

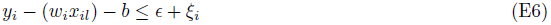

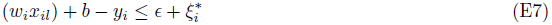

*Slack variables ξ_i_, ξ_i_* ≥ 0 *for i* = 1; 2; …; *l*; *l* = *number of data*

### 4.6 Model Selection by Leave-One-Species-Out Cross Validation

The two most popular methods for model selection in machine learning are Akaike information criterion (AIC) and Bayesian Information Criterion (BIC). These methods select models that balance model complexities and fitting, but do not directly measure predictive power. Instead, we used cross validation to measure the predictive power of how well the model performs. In cross validation, the data are divided into training and test data to separately evaluate the test data using the model learned from the training data. When k is the number of samples in k-fold cross validation, the process is called Leave-One-Out (LOO) Cross Validation. For our purposes, we adopted similar approach called Leave-One-Species-Out (LOSO) approach, meaning that we use all the data from 25 species to train the model and then test the model for the one remaining species. We repeat the cross validation 26 times, once for each species, to average the performance. The benefit of LOSO is all species will be used for training equally many times and for testing exactly once. This property also matches our practical goal of model fitting and prediction of a novel genome. The results of LOSO cross validation are illustrated in (Supplementary Figure S7) in which optimizing either the mean of the residuals or the Mean Squared Error(MSE) selected the same model.

### 4.7 Predictive Power of Our Model

We use Lasso regression and ridge regression [Smola and Schölkopf, 2004] as our baseline for comparison. The standard regression algorithms each have a MSE of approximately 580 and 17% of mean residual boundary while SVR with a one degree polynomial kernel has a MSE of 250 and 9.17% of residual mean. In other words, our model can predict the assembly performance for a new genome within approximately 9% of residual boundary. As we increase the degree of polynomial kernels, the model has less error, but risks overfitting. By LOSO cross validation, we determined SVR using RBF kernel demonstrates the best predictive power. We also evaluated the AIC for all these models, and determined that the SVR with RBF kernel has the minimum value, meaning that in obtains the best balance between model complexity and error (Supplementary Figure S7).

### 4.8 Enhanced Hybrid Error Correction

ECTools requires the low-error rate short reads be pre-assembled into high quality “unitigs” [Myers, 2005] as the basis for the long read correction. Although the input unitigs can be created by any assembler, the Celera assembler is recommended because it guarantees that every read is incorporated into a unitig, including unitigs of individual reads if necessary. This guarantee ensures that all short-read information is carried over into the produced set of unitigs and no information is lost in the preassembly process. The preassembly generates longer unitig sequences from the short-reads, creating more opportunity for alignments to span regions of high error in the long reads.

To align the preassembled unitigs to the long reads, an accurate aligner that can tolerate high local error was needed. The Mummer suite of tools provides a highly accurate nucleotide alignment script called *Nucmer* [Kurtz *et al.*, 2004]. Nucmer has many options for tuning alignments including the ability to control the maximum length of a low identity region before trimming the alignment. Because we are aligning unitigs, rather than short-reads, a consensus approach can no longer be applied to correct the long reads. Instead, the error correction selects a set of short-read unitigs that best covers each long read and uses it as a backbone for correction. Building this set is non-trivial, especially when a long read spans a repeat region. The pipeline relies on the Longest Increasing Subsequence algorithm implemented by the delta-filter program of the MUMmer suite [Salzberg *et al.*, 2012] to find the optimal set of unitigs. Once found, this unitig set can be used to correct the spanning long read. The show-snps program identifies the bases that are different between the backbone and the long read and a python script is used to make the corrections. The alignment of the unitigs to the long reads and the subsequent filtering of those alignments is quite computationally demanding so a helper script is provided so that the correction can be done on a cluster using Grid Engine.

The pipeline also implements a more conservative trimming algorithm than PacBioToCA in an attempt to preserve as much input data as possible. Firstly, the ends of the long read are trimmed if no unitig coverage is found over these bases. Internal regions without unitig coverage are considered uncorrected and are labeled at the 15% canonical PacBio error rate. Regions with unitig coverage are assumed to be corrected and assigned an error rate of 1%. The average error rate is calculated over the read using the above heuristics and compared to a user specified minimum correction identity parameter. This parameter allows the user to specify the cutoff that classifies reads as “corrected” or “uncorrected”. If a read’s average identity is below the user specified threshold, the read is split by removing the largest “uncorrected” region of the read and then recursively applying the previous steps to each resulting subreads. This algorithm helps favor sequence read continuity over pure accuracy, but because the identity parameter is user-tunable, it can easily be adjusted so identity is favored over continuity.

## Abbreviations

bp: base pair
Gbp: gigabases
Mbp: megabases
SNP: single nucleotide polymorphism

## Acknowledgements

This project was supported in part by National Science Foundation awards DBI-126383 and DBI-1350041 to MCS, and IOS-1032105 and DBI-0922738 to WRM. It was also supported in part by National Institutes of Health award R01-HG006677 to MCS. We would like to thank Adam Phillippy, Sergey Koren, and Jason Chin for their helpful discussions. We would also like to thank Susan McCouch, Lyza Maŕon, Namrata Singh, Gholson Lyon, Detlef Weigel, Panchajanya Deshpande, Senem Mavruk Eskipehlivan, Melissa Kramer, Sara Goodwin, Eric Antoniou, Cheryl Heiner, Colleen Ludka, Paul Peluso, David Rank, Pepper Weise, and Greg Khitrov for providing samples and assistance for this study.

## Contributions

H.L performed computational experiments on assembly simulation, data analysis and modeling, and wrote the manuscript. J.G developed ECTools, performed the non-simulated assemblies and wrote the manuscript. S.Y contributed to modeling and prediction using machine learning and edited manuscript. S.M performed data analysis and edited manuscript. W.R.M and M.C.S designed the study, supervised the project and wrote the manuscript.

